# Evaluation of Gene Set Enrichment Analysis (GSEA) tools highlights the value of single sample approaches over pairwise for robust biological discovery

**DOI:** 10.1101/2024.03.15.585228

**Authors:** Courtney Bull, Ryan M Byrne, Natalie C Fisher, Shania M Corry, Raheleh Amirkhah, Jessica Edwards, Lily Hillson, Mark Lawler, Aideen Ryan, Felicity Lamrock, Philip D Dunne, Sudhir B Malla

**Affiliations:** The Patrick G Johnston Centre for Cancer Research, Queen’s University Belfast, UK; School of Cancer Sciences, University of Glasgow, Glasgow, UK; Discipline of Pharmacology & Therapeutics, School of Medicine, College of Medicine, Nursing and Health Sciences, University of Galway, Ireland; Mathematical Sciences Research Centre, Queen’s University Belfast, UK; Cancer Research UK Scotland Institute, Glasgow, UK

**Keywords:** Transcriptional Signatures, GSEA, Molecular Classification, Computational Biology

## Abstract

**Background:** Gene set enrichment analysis (GSEA) tools can be used to identify biological insights from transcriptional datasets and have become an integral analysis within gene expression-based cancer studies. Over the years, additional methods of GSEA-based tools have been developed, providing the field with an ever-expanding range of options to choose from. Although several studies have compared the statistical performance of these tools, the downstream biological implications that arise when choosing between the range of pairwise or single sample forms of GSEA methods remain understudied.

**Methods:** In this study, we compare the statistical and biological interpretation of results obtained when using a variety of pre-ranking methods and options for pairwise GSEA and fast GSEA (fGSEA), alongside single sample GSEA (ssGSEA) and gene set variation analysis (GSVA). These analyses are applied to a well-established cohort of n=215 colon tumour samples, using the clinical feature of cancer recurrence status, non-relapse (NR) and relapse (R), as an initial exemplar, in conjunction with the Molecular Signatures Database “Hallmark” gene sets.

**Results:** Despite minor fluctuations in statistical performance, pairwise analysis revealed remarkably similar results when deployed using a range of gene pre-ranking methods or across a range of choices of GSEA versus fGSEA, with the same well-established prognostic signatures being consistently returned as significantly associated with relapse status. In contrast, when the same statistically significant signatures, such as Interferon Gamma Response, were assessed using ssGSEA and GSVA approaches, there was a complete absence of biological distinction between these groups (NR and R).

**Conclusions:** Data presented here highlights how pairwise methods can overgeneralise biological enrichment within a group, assigning strong statistical significance to gene sets that may be inadvertently interpreted as equating to distinct biology. Importantly, single sample approaches allow users to clearly visualise and interpret statistical significance alongside biological distinction between samples within groups-of-interest; thus, providing a more robust and reliable basis for discovery research.

## Introduction

Decreasing costs for sequencing, coupled with an increasing adoption of the FAIR principles^1^, have provided the cancer research field with a substantial amount of freely available molecular datasets derived from tumour tissue samples. To ensure that these large datasets can reveal important mechanistic insights, increased data availability has been coupled with the development of transcriptional signatures that represent important biological pathways, alongside easy-to-use algorithms that allow users to apply thousands of signatures simultaneously to these data. These are exemplified by the establishment of the Molecular Signatures Database (MSigDB)^2^ and gene set enrichment analysis (GSEA) tools^3^, providing the field with a stable set of reference templates and methods to compare across cohorts of interest. The success of these approaches has led to a rapid expansion of established signature collections in both human and mouse, most notably the MSigDB biological “Hallmark” collection^4^ and development of programming software-based GSEA tools such as the clusterProfiler^5^ and fast GSEA (fGSEA)^6^ R packages.

Given that many tumour cohorts have associated metadata linked to important features, such as clinical outcome, the application of these large collections of signatures to cohorts in conjunction with GSEA can serve as the basis for discovery and validation of biomarkers that represent the biological characteristics of the chosen features, such as prognosis. This approach is referred to as a supervised pairwise analysis, as the groups are known prior to application of the GSEA method, and these tools have been tested extensively in terms of the statistical robustness and performance in this setting^7,8^. Once identified, these biomarkers can be used as the basis for mechanistic investigations, pre-clinical model development, and/or testing of a therapeutic target.

Alongside pairwise GSEA methods, approaches for single sample methods have been developed, which differ from the pairwise approach in that they allow users to apply the same transcriptional signature collections to all samples individually in a cohort, using single sample GSEA (ssGSEA)^9^ and gene set variation analysis (GSVA)^10^. While these single sample approaches are based on different statistical models to those in pairwise analyses, the resulting outputs are based on the same gene signatures. Numerous studies have assessed the statistical robustness and performance of this range of pairwise and single sample tools separately^7,11^. Despite differences being identified between methods when assessed using statistically-driven criteria, few studies have focussed on the consequences in terms of downstream biological approaches. Given that significant pairwise GSEA results can be interpreted as representing the defining biological characteristics of a group of samples, the absence of a comparative study across all approaches means that such an interpretation may be based on incomplete evidence.

In this study, we use a fixed set of transcriptional signatures, in conjunction with a fixed clinical feature (relapse status) within a well-characterised colon cancer (CC) transcriptional cohort^12^, to perform a series of pairwise and single sample assessments in tandem. Each output is assessed based on the provided statistical values, however the primary focus of this study is to assess how representative and uniform a significant pairwise result is when assessed by single sample methods. Utilising a range of data visualisations and performance measurements, we find that statistical results from a pairwise analysis often do not align with biological distinction when using single sample outputs for the same signature. Moreover, significant signatures identified from pairwise analysis can still be poor predictive biomarkers of the clinical groups they were developed to represent.

## Methods

### Datasets

The transcriptional dataset used was previously assembled for the development of the FDA-approved stage II ColDx/GeneFx risk-of-recurrence/relapse assay, consisting of n=215 stage II primary tumours from CC patients profiled on the Almac disease-specific array, and available from ArrayExpress, accession number E-MTAB-863^12^. The cohort contained n=73 tumours from patients that went on to develop distant metastasis within 5-year of surgery to remove the primary tumour (relapse) (R) and n=142 tumours from patients that did not experience relapse within five years following surgery (non-relapse) (NR). The E-MTAB-863 CEL files were imported into Partek Genomics Suite (PGS; v6.6) and RMA normalised then log2 transformed. The probesets on the array were collapsed by importing the normalised data into R (v3.3.2 or later) and, using the ‘*collapseRows*’ function from WGCNA (Weighted Gene Coexpression Network Analysis, RRID:SCR_003302) package (v1.61)^13^, selecting the probeset with the highest mean expression per gene.

### Differential gene expression analysis

Differential expression analysis (DEA) was performed to measure differentially expressed genes between R and NR CC. DEA was performed using the *limma* R package (v3.54.2). Following DEA, genes were ranked using three different metrics, 1) the *t*-statistic (*t*-stat), 2) the Log Fold Change (LogFC), and 3) the combination of LogFC and p-value (LogFC*-Log10(p-value); hereafter as “combined”). The addition of p-value to LogFC adds statistical significance to the directionality of LogFC. Separately, DEA was also performed for another comparison between tumours classified as PDS1 and PDS3, using the *PDSclassifier* package^14^ with resulting groups being assessed using the same metrics and thresholds applied to the R/NR analyses.

### Pairwise analysis

To perform pairwise analysis two R packages were used, *clusterProfiler* (v4.6.2) and *fgsea* (v1.24.0) and a random seed of 127 was set. Biological pathways were investigated using the Hallmark gene sets from the MSigDB accessed through the *msigdbr* package (v7.5.1). Pre-ranked GSEA was first performed using the GSEA function in *clusterProfiler* with 1000 permutations (nPermSimple = 1000, minGSSize = 1, maxGSSize = Inf). Enrichment plots for GSEA were produced using the *gseaplot2* function in the *enrichplot* R package (v1.18.4). GSEA was next conducted using the *fgsea* R package with the same parameters as *clusterProfiler* (nPermSimple = 1000, minSize = 1, maxSize = Inf). Enrichment plots of *fGSEA* were produced using the *plotEnrichment* function from the *fgsea* package. The online tool, GenePattern^15^, https://cloud.genepattern.org, was also used to perform a pre-ranked pairwise analysis, GSEAPreranked (v7.4.0). The Hallmark gene set collection was selected, ‘h.all.v2023.2.Hs.symbols.gmt’. Default parameters were set except for ‘collapse dataset’ which was set to ‘FALSE’. Normalised enrichment score (NES) and false discovery rate (FDR) values were recorded for each gene set within the two groups (R vs NR; PDS1 vs PDS3). A gene set with an FDR *q*-value below 0.05 was deemed significant.

### Single sample analysis

To perform single sample analysis the R/Bioconductor package *GSVA* (v1.46.0) was used which facilitates ssGSEA^9^ and GSVA^10^. ssGSEA was performed with Hallmark^4^ gene sets from MSigDB^2^ and method set to “ssgsea”. GSVA was performed with Hallmark gene sets from MSigDB and the default parameters.

### Single sample analysis heatmaps

For both ssGSEA and GSVA, matrix was formatted to include only Interferon Alpha Response, Interferon Gamma Response and Epithelial Mesenchymal Transition (EMT), as previously identified to be most significant by GSEA. The single sample scores were converted to Z-scores and were plotted using the *ComplexHeatmap* (v2.14.0) R package and were grouped using their respective groups (R vs NR; PDS1 vs PDS3).

### Data visualisation

Additional visualisation R packages used for single sample analysis included: *smplot2* (v 0.1.0), *ggridges* (v 0.5.4), *easyGgplot2* (v 1.0.0.9000), *pROC* (v 1.18.5), *randomForest* (v 4.7 -1.1) and, *waterfalls* (v 1.0.0).

### Statistics

The statistical report was generated on RStudio (4.2.2). A Student’s *t*-test, from the *stats* (v 4.2.2) R package, was used to calculate significance of single sample scores between groups (NR compared to R and PDS1 compared to PDS3). The *cor.test* function from the *stats* (v 4.2.2) R package, with “pearson” method selected, was used for correlation analysis between single sample enrichment scores for selected significant gene sets. The *cutpointr* function in the *cutpointr* (v 1.1.2) R package was used to find the optimal cutpoint for the single sample scores. Once calculated the single sample scores were centred around the cutpoint resulting in a stratification of high and low scores for each of the gene sets being tested.

### “dualgsea”

The pairwise method, fGSEA^16^ and single sample method, ssGSEA^9^ have been combined to create an open source R-based function named “*dualgsea*”, https://github.com/MolecularPathologyLab/Bull-et-al. The function enables the user to apply the above statistical analysis and visualisations between two groups-of-interest.

## Results

### Variations in differential gene expression outputs across a range of methods do not alter overall GSEA results

A typical goal when analysing bulk transcriptomic data, is the identification of discriminatory biological signalling cascades that can serve as biomarkers to distinguish between group(s)-of-interest; an output that can rapidly be delivered using transcriptional signatures in conjunction with *in silico* analytical tools, such as pairwise gene set enrichment analysis (GSEA)^3^ (Figure 1A). The initial step in this GSEA process requires all genes in the expression matrix to be ranked based on their differential expression between the groups-of-interest. For example, when using *limma*^17^ for microarray or *DESeq2*^18^ for RNA-seq, a ranked list of genes can be produced based on *t*-statistics (*t*-stat) or Log Fold Change (LogFC) values, both of which also provide directionality (up/down) according to the groups used. To assess the outputs from each ranking metric, we compared the ranked order of genes following the application of three approaches based on: 1) *t*-stat, 2) LogFC, and 3) the combination of LogFC and p-value (LogFC * -Log_10_(p-value); hereafter stated as combined) on expression profiles from n=15,723 genes derived from n=215 FFPE stage II colon cancer samples (E-MTAB-863)^12^, where patients whose cancer relapsed following surgery (n=73) compared to those who did not (NR; n=142) was used as an exemplar pairwise GSEA comparison (Figure 1B). Considering only the top and bottom 100 genes ordered based on *t*-stat (0.6% of genes overall), gene ordering based on LogFC, or the combined rank, remained remarkably stable. The top/bottom ranked genes identified using each method remain highly enriched at the extremes relative to *t*-stat ranking (Figure 1B). When the genes were ranked by logFC the majority (86%) of the top 100 genes fell within the top 500 genes when ranked by *t*-stat and the remaining were represented within the top 2,707 genes. With the combined rank, 100% of the top 100 genes were represented within the top 300 genes when ranked by *t*-stat.

**Figure 1.**
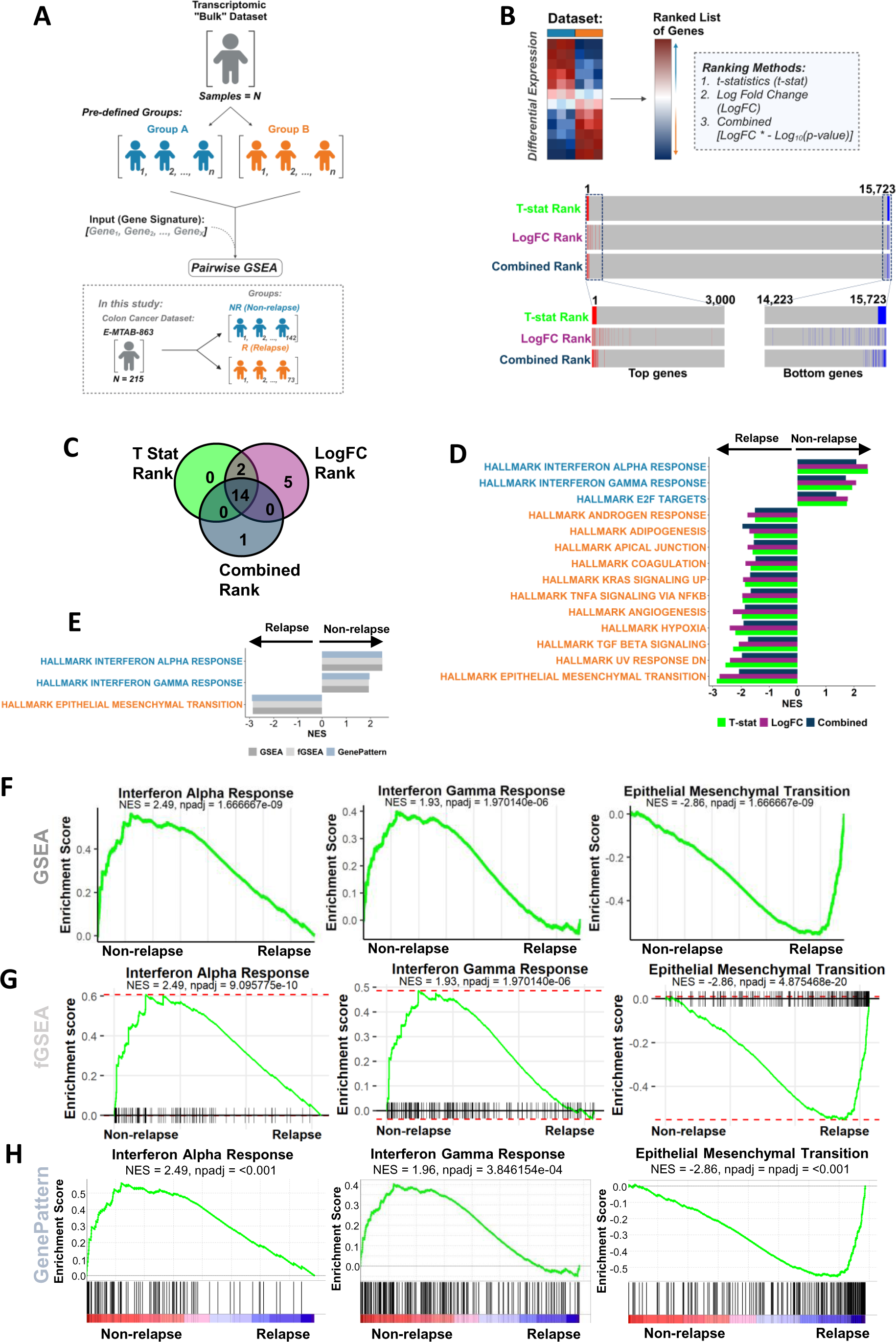
Differential gene expression analysis and pairwise analysis of the discovery cohort. (A) Schematic of the differential expression analysis and pairwise analysis. (B) Workflow of differential expression analysis and ranked position of the top 100 differentially expressed genes and bottom 100 genes in NR when ranked by *t*-stat and the position of these genes when ranked by logFC and combined. (C) Venn diagram of the significant Hallmark signatures (padj < 0.05) from GSEA when genes were ranked by *t*-stat, logFC, and combined. (D) Significant Hallmark signatures (padj < 0.05) identified from clusterProfiler GSEA when genes were ranked by *t*-stat, logFC, and logFC combined with the p-value ordered by NES. (E) clusterProfiler GSEA, fGSEA, and GenePattern pre-ranked GSEA of the significant Hallmark gene sets. (F-H) clusterProfiler GSEA (F), fGSEA (G), GenePattern (H) comparing NR CC (n=142) to R CC (n=73) for Interferon Alpha Response, Interferon Gamma Response and EMT.

To test if there were more profound downstream consequences of these small pre-ranking gene order fluctuations, GSEA in clusterProfiler was performed^5^ using each of these ranking metrics on the n=50 MSigDB ‘Hallmark’ gene sets. These analyses revealed that all three ranking methods resulted in remarkably consistent gene sets being returned as significant (FDR adjusted p-value < 0.05; *t*-stat =16/50, LogFC = 21/50, combined = 15/50), n=14 of the n=22 total significant gene sets identified as common across from all three ranking methods (Figure 1C; Supplementary Figure 1A). When the normalised enrichment score (NES) is assessed to measure directionality, the direction of the n=14 overlapping significant gene sets identified remained entirely consistent (Figure 1D), meaning that regardless of the pre-ranking method used for these GSEA analyses, the biological interpretation will remain the same. Furthermore, when gene sets that were identified as significant by one method but not by the others, these were all enriched with the same directionality yet just below the statistical significance threshold: again, confirming the similarities in outputs for GSEA using all three pre-ranking methods (Supplementary Figure 1A).

### Pairwise GSEA methods provide results with consistent downstream interpretation

As there were minimal differences in the GSEA outcome with the three ranking methods, *t*-stat was used for the remainder of this study. Since the introduction of the original GSEA method, several updated methodologies have been developed and in this study we examined three derivatives of the GSEA method: 1) fast GSEA (fGSEA)^6^, 2) GSEA via clusterProfiler^5^ (as used in Figure 1), which are both R-based tools, and 3) GSEA^3^ from the Broad Institute GenePattern^15^ Server. The GSEA tool from GenePattern performs standard GSEA with default signal-to-noise for ranking genes, however, the server also provides users with a separate module called ‘GSEAPreranked’, where users can provide their own pre-ranked gene list prior to analysis. To test outputs from each of these GSEA methods, relapse (R) (n=73) and non-relapse (NR) (n=142) groups were compared across the CC cohort previously used (E-MTAB-863), where these methods consistently identify the same common statistically significant gene sets as identified in Figure 1E, additionally the directionality of the NES for gene sets is consistent (Supplementary Figure 1B). Between these three methods, n=3 gene sets were consistently upregulated in the NR group, including Interferon Alpha Response and Interferon Gamma Response, and n=11 gene sets were upregulated in the R group, such as EMT (Figure 1F-H); gene sets that have previously been associated with prognosis in multiple cancer types, including colorectal cancer^19, 20^.

### Single sample GSEA methods provide biological insights that may be masked when using pairwise GSEA alone

Single sample GSEA (ssGSEA)^9^ has been proposed as an extension of the GSEA method, one which can provide signature enrichment scores for each individual sample, rather than the summarised “average” scores within groups of samples provided by pairwise GSEA, making it suitable for both biological discovery and post-hoc assessments of individual samples within any established groups-of-interest ^21^ ^22^. Therefore, to compare the results obtained from GSEA (Figure 1) with those from the single sample approaches, we explored two such methods: 1) ssGSEA^9^, and 2) gene set variation analysis (GSVA)^10^ within our discovery cohort (Figure 2A). Using the top three significant gene sets identified in Figure 1E, namely Interferon Alpha Response, Interferon Gamma Response and EMT, these single sample approaches were run using the *GSVA* R package by selecting either the “ssGSEA” or “gsva” method. A correlative analysis was performed between the resulting ssGSEA and GSVA scores, which revealed that both single sample methods were highly correlated, with a significantly positive correlation across all three gene sets (R>0.8, p<0.0001; Figure 2B). These results suggest that while the algorithms are different, the output of either single sample methods provide consistent results.

**Figure 2.**
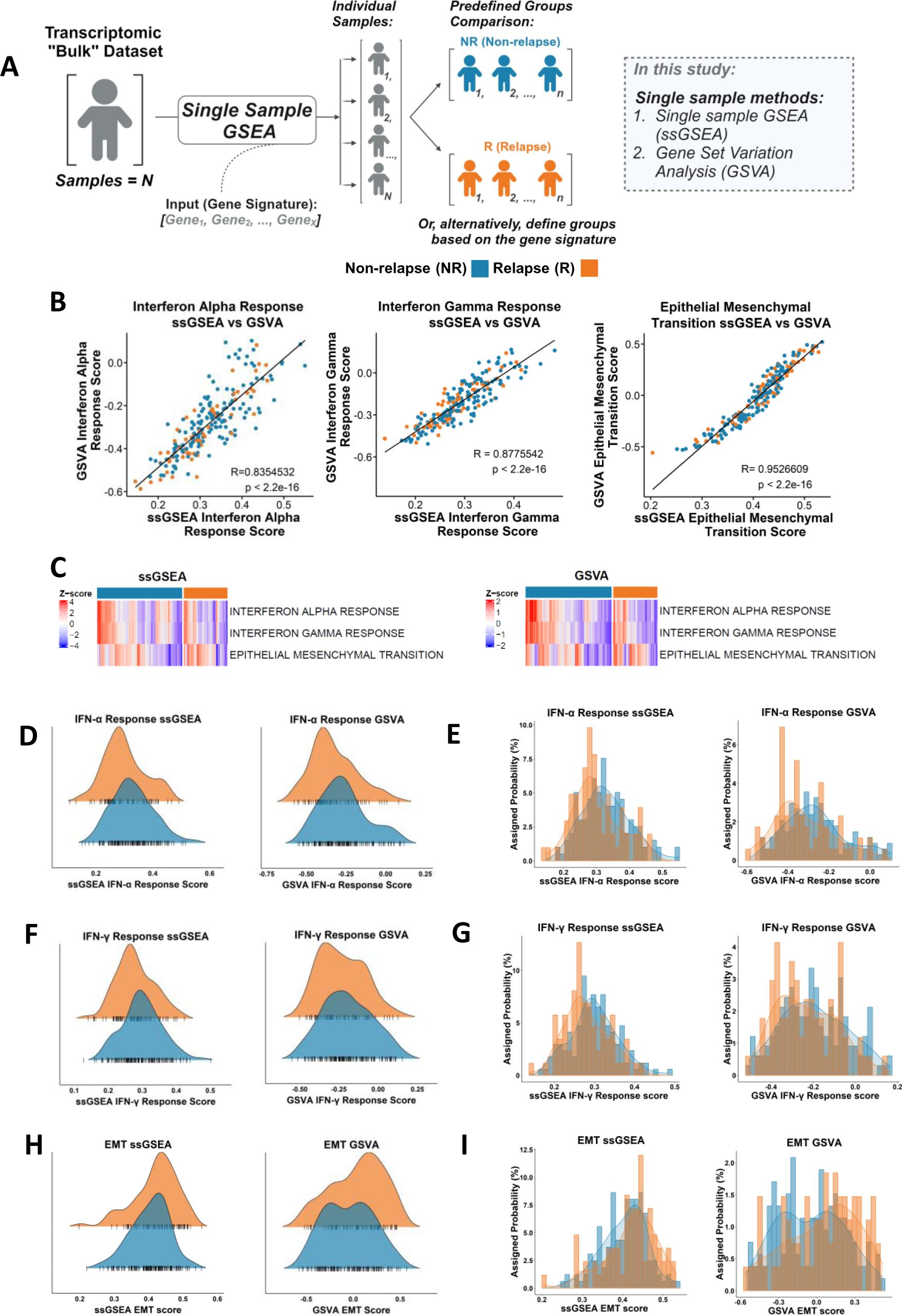
Comparison of the single sample methods, ssGSEA and GSVA. (A) Schematic of standard single sample analysis workflow. (B) Scatterplot showing the correlation of ssGSEA scores and GSVA sores for the Hallmark Interferon Alpha Response (Pearson correlation coefficient, r = 0.835), Interferon Gamma Response (r = 0.878) and EMT (r = 0.953). (C) Heatmap of ssGSEA and GSVA scores for Interferon Alpha Response, Interferon Gamma Response and EMT comparing NR and R. (D - I) Distribution of ssGSEA and GSVA scores. (D and E) Distribution of the ssGSEA and GSVA scores for the Interferon Alpha Response signature in the R (orange) and NR (blue) samples depicted using kernel density plots (D) and histograms (E). (F and G) Distribution of the ssGSEA and GSVA scores for the Interferon Gamma Response signature in the R (orange) and NR (blue) samples depicted using kernel density plots (F) and histograms (G). (H and I) Distribution of the ssGSEA and GSVA scores for the EMT signature in the R (orange) and NR (blue) samples depicted using kernel density plots (H) and histograms (I)

Assessment of the ssGSEA and GSVA scores for the three gene sets that were significantly different between the NR and R groups using GSEA, namely Interferon Alpha Response and Interferon Gamma Response and EMT, revealed that there were comparable quantities of high and low expression samples in each group, as indicated by the blue-to-red colours in the heatmap (Figure 2C). To test this, a series of quantitative assessments were performed using scores for the significant signatures using GSEA. Although the two clinical groups may appear statistically significant for these single sample scores (Supplementary Figure 2C-H), both clinical groups fall under the same distribution scale (Figure 2D-I), thus implying in biological terms, they are not distinct for the signatures, which contradicts with GSEA output. The range of ssGSEA scores showed large overlap between R and NR samples, Interferon Alpha Response had 95.3% overlap between R and NR, Interferon Gamma Response had 97.7% overlap between R and NR and EMT had 99.1% overlap between R and NR. With respect to the GSVA results, Interferon Alpha Response scores had 95.8% overlap between R and NR, Interferon Gamma Response had 98.1% overlap and EMT had 98.6% between R and NR. Overall, these data highlight how even the most statistically significant pairwise GSEA results may not be sufficient to identify transcriptional signalling that is discriminatory between samples across two tumour groups.

### Visualisation of ssGSEA score is essential to ensure that statistical significance between sample groups also represents distinct biology

There are a range of biomarker performance metrics that can be used to objectively test and enumerate how well individual signatures represent the signalling within different groups of samples. Therefore, a series of analyses were conducted to test the predictive value of the most significant signatures identified by pairwise GSEA approaches (n=3) in identifying the specific groups-of-interest that they were enriched in. We performed receiver operating characteristic (ROC) analysis with the ssGSEA/GSVA scores and examined the area under curve (AUC). NR patients displayed statistically significant enrichment in Interferon Alpha and Interferon Gamma Response, implying that these signatures are contributing factors to favourable outcome in NR patients (Supplementary Figure 2C-E), albeit GSVA Interferon Gamma Response did not show any statistically significant enrichment in the NR samples (Supplementary Figure 2F). However, if both interferon response signatures were then to be used to develop a risk stratification tool to predict patient relapse status, the models developed based on these signatures would perform underwhelmingly with the AUC approximately ranging between 0.57 – 0.62 (Figure 3A, 3C). Furthermore, although there are more NR (n=142) than R cases (n=73), when stratified into high and low groups for the Interferon Alpha and Interferon Gamma Response signature scores using both ssGSEA and GSVA, based on the optimal cut-offs defined by the AUROC analyses, ∼30-50% of relapse patients have high Interferon Alpha and Interferon Gamma Response scores (Figure 3B, 3D). Likewise, regardless of its statistical significance (Supplementary Figure 2G-H), the EMT ssGSEA and GSVA scores also perform poorly (AUC 0.60), with low sensitivity and specificity as a relapse-specific biological signature for the purpose of risk stratification (Figure 3E-F).

**Figure 3.**
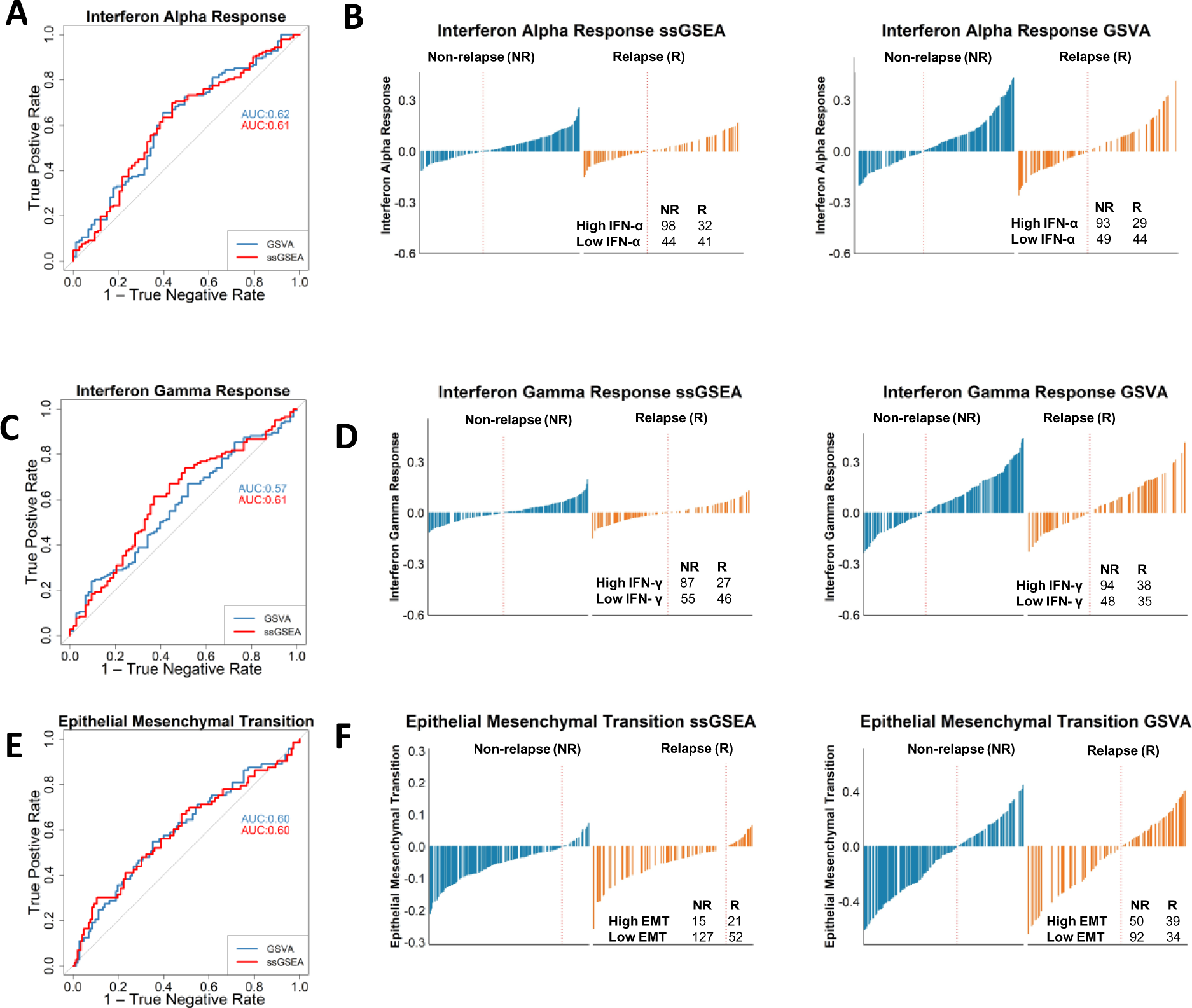
Application of single sample analysis as a predictor for relapse. (A) ROC curve using Interferon alpha response ssGSEA and GSVA scores to predict NR had an AUC ranging between 0.61 – 0.62. True positive rate is when the sample is classified as high Interferon Alpha Response, and the case was a NR. The true negative rate is the proportion of true negatives, when a sample is a NR cases without high Interferon Alpha Response. (B) Interferon Alpha Response waterfall plots show when stratified into high and low groups for the Interferon Alpha Response with both ssGSEA and GSVA scores, we found a greater number of NR patients classed as high (n=98 [75.4%], n=92 [76.0%]) respectively compared to R (n=32 [24.6%], n=29 [24.0%]) respectively. (C) Interferon Gamma Response ROC AUC values ranging between 0.57 – 0.61. True positive rate is when the sample is classified as high Interferon Gamma Response, and the case was a NR. The true negative rate is the proportion of true negatives, when a sample is a NR cases with a low Interferon Gamma Response score. (D) Interferon Gamma Response waterfall plots show when stratified into high and low groups for the Interferon Gamma Response with both ssGSEA and GSVA scores, there were greater number of NR patients classed as high (n=86 [76.1%],; n= 95 [71.4%] respectively compared to R (n=27 [23.9%],; n=38 [28.6%] respectively (E) EMT ROC AUC values of 0.60. True positive rate is when the sample is classified as high EMT, and the case was a relapse. The true negative rate is the proportion of true negatives, when a sample is a NR case with a low EMT score. (F) EMT waterfall plots show when stratified into high and low for EMT for ssGSEA we found that a greater number of R patients classified as high (n=22 [59.5%], compared to NR (n=15 40.5%). GSVA found a higher number of NR patients with a high EMT score (n=50 56.2%) compared to R (n=39 43.8%)

Taken together, while each of these three signatures have been repeatedly shown to provide statistical significance in terms of association with relapse outcomes, this is primarily due to small (albeit statistically significant) differences in sample distributions, meaning that the biological signalling these signatures are based on cannot be interpreted as reflecting distinct mechanistic phenotypes or biological cascades between the two groups-of-interest.

### Pathway-derived subtype serves as an exemplar for performing biological discovery using a single sample approach

As shown above, pairwise methods comparing relapse and non-relapse tumours can provide users with statistically significant results, however these clinically distinct groups do not represent uniformly biological distinct transcriptional subtypes. Therefore, to test the performance of pairwise and single samples GSEA methodologies in groups of samples that represent biologically distinct entities, we next performed these analyses contrasting tumours based on our recent pathway-derived subtypes (PDS) ^14^ which identified three statistically and biologically distinct subtypes; PDS1-3.

In this current study we now segregate our transcriptional cohort into these three PDS classes (this dataset was not used in the original study) and perform a series of GSEA/ssGSEA assessments on PDS1 (characterised by high MYC signalling) and PDS3 (characterised by low MYC signalling) in conjunction with the performance metrics and visualisations used so far (Figure 4A). Comparative analysis using the Hallmark gene sets collection and pairwise GSEA, similar to the relapse-based comparisons, highlights a highly significant statistical difference between PDS1 and PDS3 for MYC Targets V1 gene set (hereafter MYC V1; Figure 4B). Importantly, unlike the assessment on R versus NR in the same cohort (Figure 1-3), comparison of PDS1 to PDS3 clearly shows both statistical significance and biological distinction when using single sample approaches (Figure 4C). Most importantly, unlike our earlier analyses based on GSEA results comparing R and NR samples, these new assessments across a known biology, reveal a remarkable difference and minimal overlapping distribution for MYC V1 ssGSEA score, with only 6.7% of ssGSEA scores overlapping between PDS1 and PDS3 (Figure 4D-E), implying that PDS1 and PDS3 can be considered as representing truly distinct biological groups for MYC V1. This is further confirmed using ROC analysis, from both ssGSEA and GSVA MYC V1 scores, which proves a sample will be classified as high MYC V1 when the sample is PDS1 with an AUROC = 0.99 (Figure 4F-G). We have created an open source parallel pairwise/single sample R-based function “*dualgsea*”, https://github.com/MolecularPathologyLab/Bull-et-al. The function produces multiple visualisations and statistical analysis options that enables users to perform a broad characterisation of their samples and groups-of-interest (Figure 4H).

**Figure 4.**
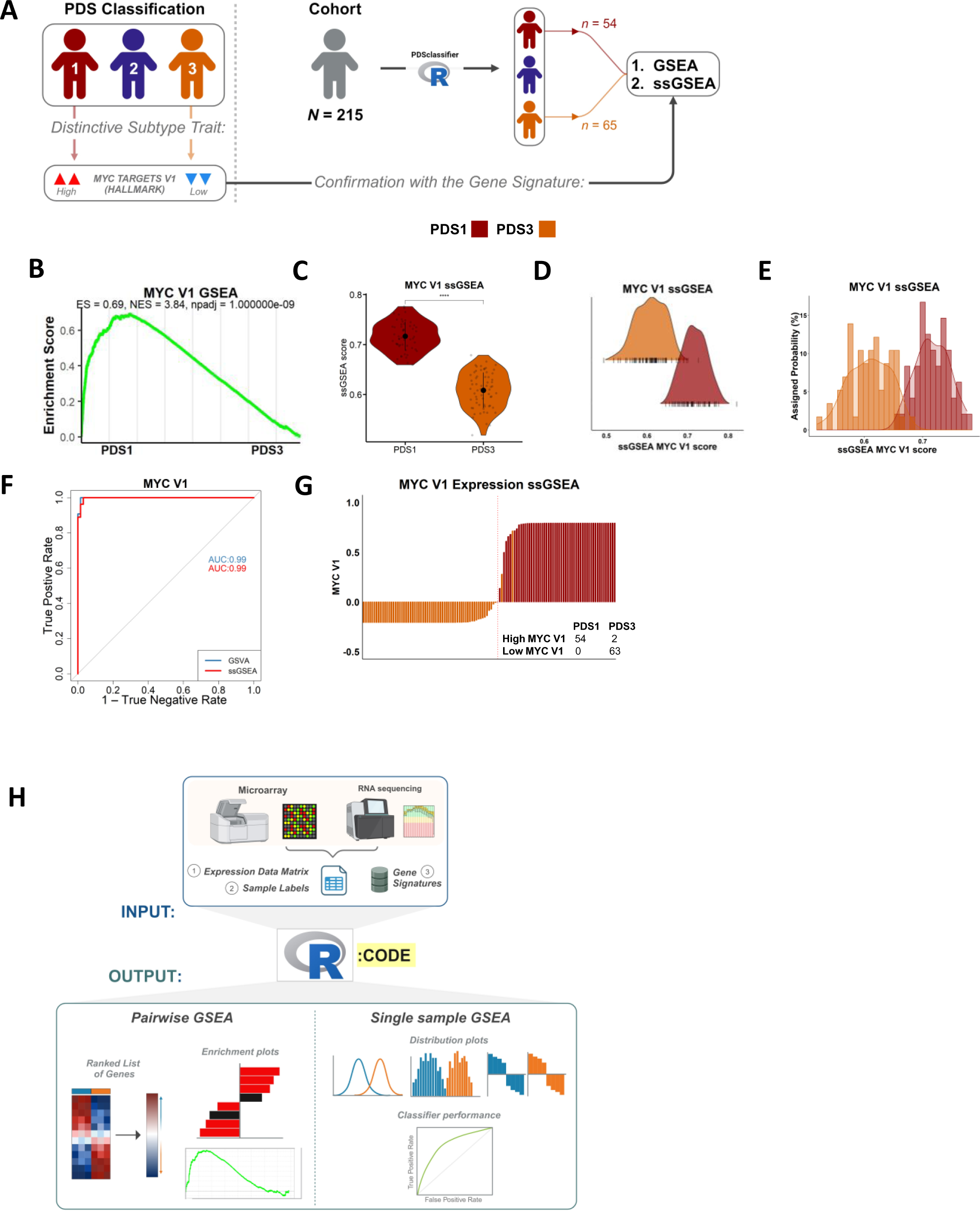
The use of single sample analysis provides distinct biology between groups. (A) Schematic of application of pathway analysis methods when applied to PDS classification. (B) GSEA revealed MYC targets V1 is enriched in the PDS1 group compared to the PDS3 group. (C) ssGSEA scores show significant difference of MYC targets V1 expression between PDS1 and PDS3 groups (**** p-value < 0.0001). (D & E) ssGSEA scores for PDS1 and PDS3 show little overlap of MYC targets V1 expression between groups. (F) ROC curve shows that the MYC V1 scores enable discrimination between PDS1 and PDS3. AUC value of 0.99. True positive rate is when the sample is classified as high MYC V1, and the case was PDS1. The true negative rate is the proportion of true negatives, when a sample is a PDS1 without high MYC V1. (G) Stratification of MYC V1 high and MYC V1 low ssGSEA scores showed that PDS1 was classified as high MYC V1 (n=54 [96.4%] and PDS3 contained only samples with a low MYC V1 score (n=63 [100%]).

## Discussion

In this study, we initially set out to provide a comparison of a number of well-established gene set enrichment analysis (GSEA) methods, with particular emphasis on how choices of standard bioinformatic pipelines can lead to differences in downstream biological interpretation. As an exemplar of this, we assessed how consistent a significant pairwise GSEA result is between pairwise approaches and also when the same signature is assessed using single sample GSEA methods. These analyses highlight concordance *within* pairwise or single sample approaches, however despite similar statistical performance, data presented here provides a clear indication for how vastly different downstream interpretation of results can be derived when using pairwise or single sample methods for the same transcriptional signatures. Pairwise methods provide the user with strong statistical-based evidence of differences in signature expression between two selected groups of samples, however this can result in confusion when interpreting the biological significance of these differences, as illustrated by enrichment scores across individual samples strongly overlapping between and within groups. These results strongly support the use of single sample methods for class discovery and mechanistic biomarker development/testing, given their consistency and robustness in identifying distinct biological signalling between defined groups of samples. Many previous studies have focussed on the statistical advantages and limitations of GSEA methods, providing the field with important information on performance metrics for each algorithm^7^. While these algorithms were developed to identify ***statistical*** significance between user-selected groups of samples, they can occasionally be interpreted as representing ***biologically*** distinct groups; a point that becomes even more important if the results from GSEA-based methods are used to guide development of new pre-clinical models that are interpreted as faithfully representing the clinical group-of-interest, or used as the basis of developing prognostic/predictive biomarkers to guide clinical decision-making.

Data presented in this paper does not challenge the importance of studies using GSEA methods, as we clearly demonstrate their value in identifying robust statistically distinct groups. Our current study aims to provide an example of the consequence of method selection for biological end-users with a primary interest in using these tools to identify biologically distinct mechanistic signalling between two groups. For such end-users, we propose that emphasis should be placed on more widespread use of visualisation methods at an individual sample resolution, rather than the use of statistical values alone, to ensure there is a clear distinction between the groups being compared^23^. This point is particularly important for biomarker discovery, where there is a requirement for the most robust and discriminatory features that can be used to predict tumour groups with high sensitivity and specificity. In addition, the identification of representative biological cascades that are both statistically significant and biologically distinct between the two groups across a cohort of tumours is increasingly important in the era of precision medicine, where interrogation of transcriptional data can be used as the basis for development and testing of subtype-specific therapeutic targets aimed at these patient groups.

An important feature for performing pairwise GSEA is the ranking of differentially expressed genes. Our analyses highlight that the positions of individual differentially expressed genes in an overall list will vary when using different ranking options. These results provide a clear example of how the use of some of the most widely accepted tools for differential gene expression analyses can lead to different users identifying conflicting biomarkers for the same phenotypes in the exact same datasets. However, we find that the effects on using different pre-ranking methods to rank genes for pairwise approaches have minimal effects on biological interpretation when using downstream pathway analyses with any GSEA method. As such, these data again support the use of pathway-level gene signatures as a more representative way of measuring true biological phenotypes in transcriptional data, over the use of individual gene-level biomarkers that can be undermined by technical biases inherent in method choices for gene ranking. This single sample approach was used as basis for class discovery within our recent pathway-derived subtypes (PDS) ^14^ study, which used ssGSEA scores to identify three biologically distinct classes of colorectal cancer that was found to have prognostic value.

The cancer research field is accustomed to the heavy reliance on statistical thresholds as the primary criteria for significance, as they provide users with a quantitative reference in support of their findings. In data presented here we clearly show that additional visualisation of these same data can lead to questions over the true biological significance of such results. In this setting, if GSEA tools were used for discovery, the biological signalling used as the basis for mechanistic studies could be indistinguishable across samples from these different clinical groups, despite such signalling being based on statistically sound evidence. Moving forward, it is essential to find a balance between statistical significance and biological relevance, utilising visualisation techniques and analysis methods, including distribution plots and ROC curves, to validate and contextualise findings. To ensure users can recapitulate the approaches used here, we have developed an open source parallel pairwise/single sample R-based function, “*dualgsea*” https://github.com/MolecularPathologyLab/Bull-et-al, which provides multiple data visualisation outputs and statistical tests, enabling all users to perform a comprehensive assessment of their samples and groups-of-interest as shown in the comparison of PDS1 vs PDS3 (Figure 4H).

Overall, our study sheds new light on the nuances between established gene set enrichment methods, highlighting the challenges in interpreting results across different methods. The work presented illustrates how a highly significant pairwise result does not always translate to a significant single sample result when the same transcriptional data is analysed using the same gene signatures. By carefully navigating these methods and their implications, researchers can uncover novel meaningful biological insights from transcriptional data.

## Supporting information

Bull et al., GSEA Supp Figures

## Acknowledgements

This work was supported by a CRUK early detection grant (A29834), a CRUK International accelerator programme, ACRCelerate, (A26825), a UK Medical Research Council (MRC) National Mouse Genetics Network programme (MC_PC_21042)

## Data Availability Statement

Data is available in a public, open access repository. The “*dualgsea*” scripts used in this current study are publicly available at https://github.com/MolecularPathologyLab/Bull-et-al.

## Supplementary Figure Legends

**Supplementary figure 1. Ranking metrics of differentially expressed genes for GSEA have little impact on GSEA results and GSEA methods have little variation.** (A) 50 Hallmark gene sets from clusterProfiler GSEA when genes were ranked by *t*-stat, logFC, and combined, highlighting the significant (padj < 0.05) hallmarks that are associated with all three ranking methods. (B) clusterProfiler GSEA, fGSEA, and GenePattern pre-ranked GSEA 50 Hallmark gene sets ranked by *t*-stat.

**Supplementary figure 2. Comparison of single sample analysis methods.** (A) ssGSEA heatmap of 50 Hallmark gene sets. (B) GSVA heatmap of 50 Hallmark gene sets. (C) Significance between NR and R ssGSEA scores for Interferon Alpha Response (** p <0.01) (D) Significance between NR and R GSVA scores for Interferon Alpha Response (** p <0.01). (E) Significance between NR and R ssGSEA scores for Interferon Gamma Response (* p <0.05). (F) No significance between NR and R GSVA scores for Interferon Gamma Response (ns). (G) No significance between NR and R ssGSEA scores for EMT (ns) (H) Significance between NR and R GSVA scores for EMT (* p <0.05).

