## Supplementary figures and images for "Evaluation of Gene Set Enrichment Analysis (GSEA) tools highlights the value of single sample approaches over pairwise for robust biological discovery"

### Bull et al., GSEA Supp Figures

#### Supplementary figure 1

A

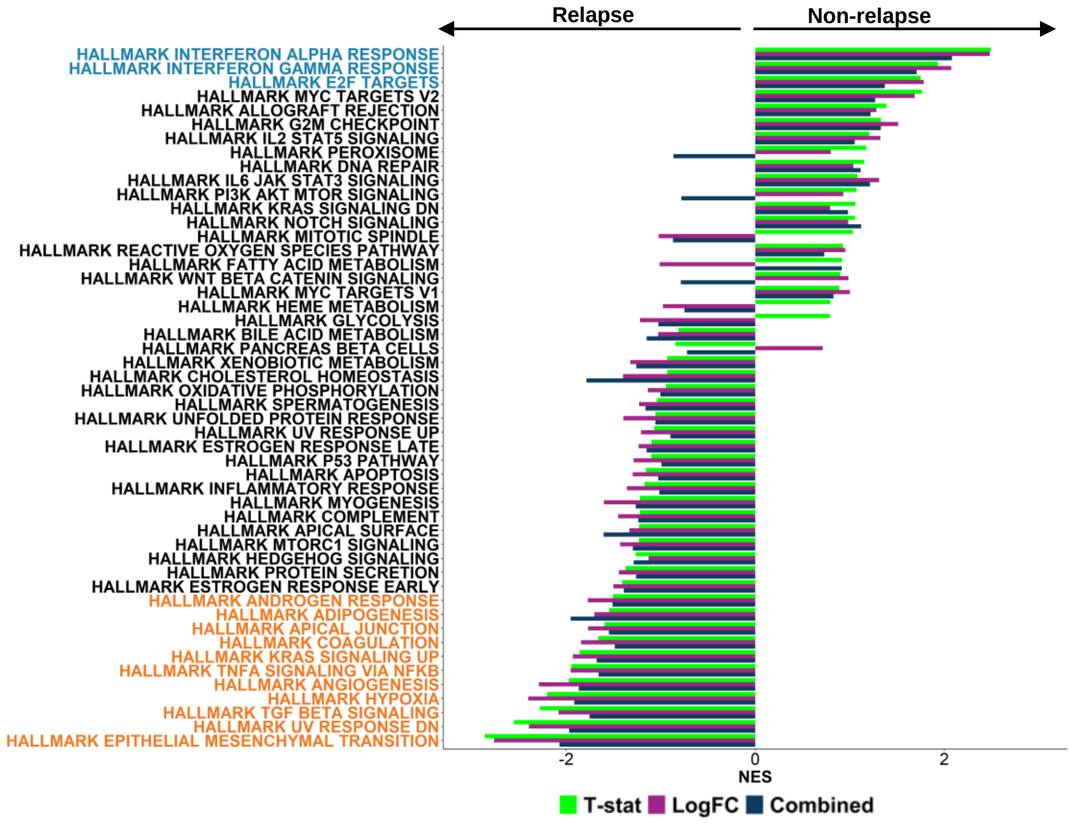


B

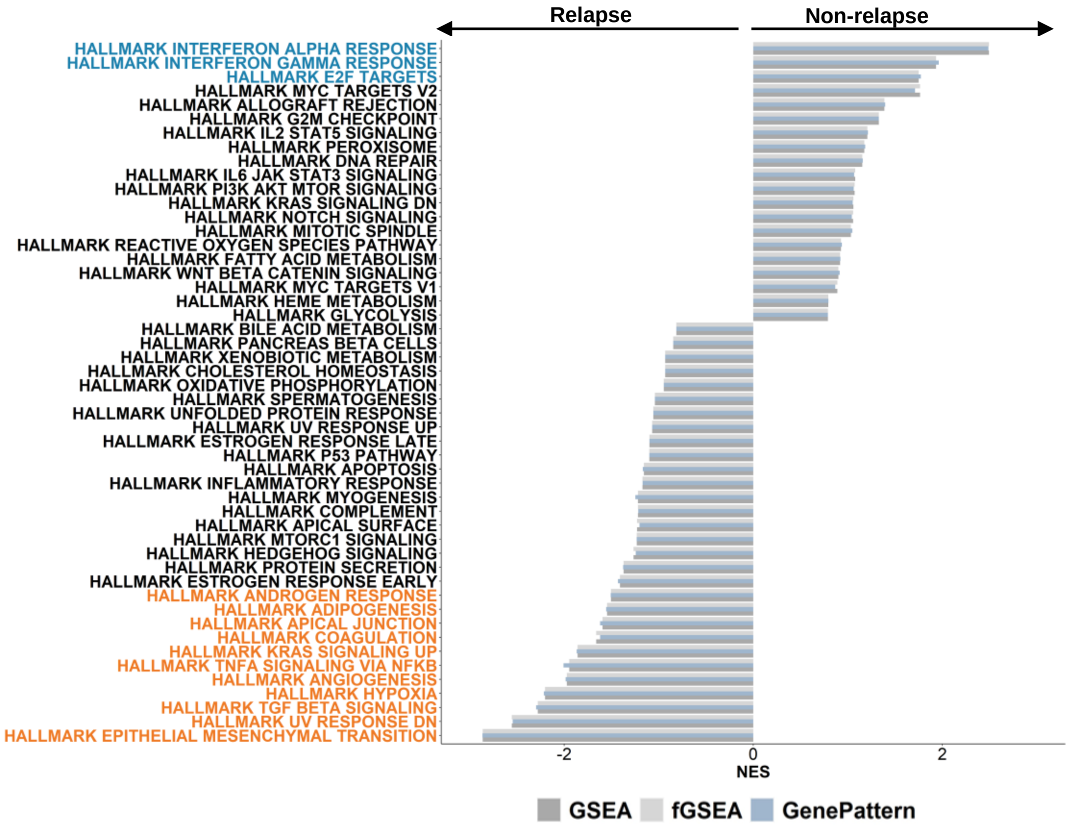

#### Supplementary figure 2

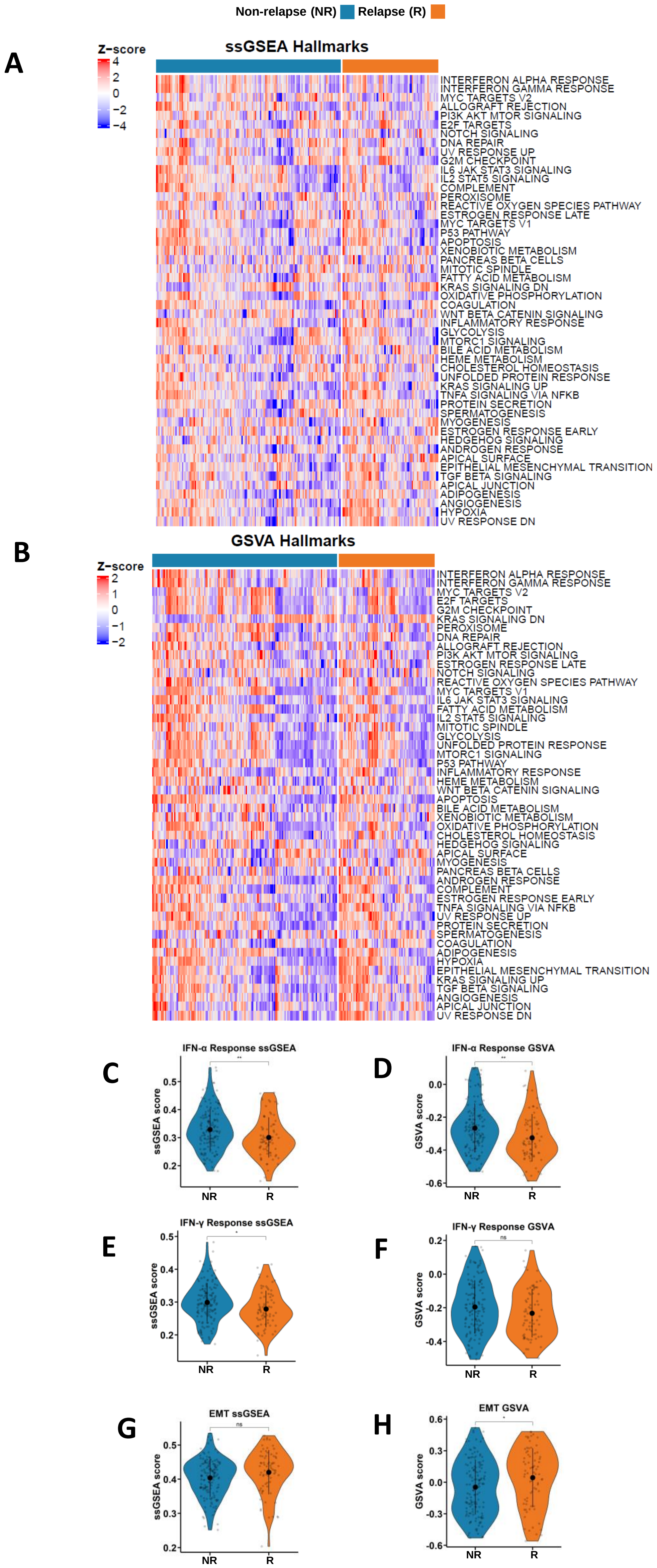
